# Parameters that influence bipartite reporter system expression in *C. elegans*

**DOI:** 10.1101/2024.03.01.583016

**Authors:** Emma Knoebel, Anna Brinck, Michael L. Nonet

## Abstract

The development of bipartite reporter systems in *C. elegans* has lagged by more than a decade behind its adoption in Drosophila, the other invertebrate model commonly used to dissect biological mechanisms. Here, we characterize many parameters that influence expression in recently developed *C. elegans* bipartite systems. We examine how DNA binding site number and spacing influence expression and characterize how these expression parameters vary in distinct tissue types. Furthermore, we examine how both basal promoters and 3’ UTR influence the specificity and level of expression. These studies provide both a framework for the rational design of driver and reporter transgenes as well as molecular and genetic tools for the creation, characterization, and optimization of bipartite system components for expression in other cell types.

## Introduction

Transgenic animals are a powerful component of the toolbox for the study of cellular and molecular processes in *C. elegans.* Transgenes enable performing a variety of experiments that are not feasible using wild type animals. These include examining the tissue specificity of gene function (e.g. Mahoney et al., 2008), assaying the temporal requirements of gene activity (e.g. Zhang et al., 2015), visualizing developmental events (e.g. Chai et al., 2012) and subcellular processes *in vivo* over time (e.g. Nadiminti and Koushika, 2022), monitoring the concentration and localization of cellular metabolites over time (e.g. Venkatachalam et al., 2016) and selective labeling for biochemical studies (e.g. Artan et al., 2021). Hence fostering the ability to create these tools promotes innovation in research by expanding the approaches that can be harnessed to dissect specific cellular and molecular mechanisms.

Techniques for building transgenic *C. elegans* animals have developed rapidly in recent years as novel methods that facilitate single copy insertion of moderate sized DNA constructs have come on board (Nance and Frøkjær-Jensen, 2019). These include transposon mediated insertion (Frøkjær-Jensen et al., 2014), insertions mediated by repair of double stranded breaks (Frokjaer-Jensen et al., 2008; Dickinson et al., 2013; Arribere et al., 2014; Paix et al., 2014; Silva-García et al., 2019; El Mouridi et al., 2022; Stevenson et al., 2023) and the use of site specific recombinases to catalyze insertions (Nonet, 2020; Yang et al., 2022; Nonet, 2023). Techniques that facilitate the creation of multi-copy insertions have also been developed based on generating and repairing double stranded breaks at either specific or random positions (Yoshina et al., 2016; Noma and Jin, 2018; El Mouridi et al., 2022). Although these methods have greatly improved the landscape for creating transgenic animals, several limitations still often cripple the ability to create robust transgenic tools.

One aspect of transgenesis in *C. elegans* that remains problematic is creating transgenic animals that express the desired product at the <u>appropriate</u> level. One common desired outcome is a transgenic line that expresses at very high levels to facilitate tracking/analysis of animals carrying the transgene. Single copy insertions are the simplest stable transgenes to create and are molecularly well-defined, but often express the desired product at sub-optimal levels. Multi-copy insertions often yield higher level expression, but have major drawbacks. They are generally difficult, if not impossible, to create with a specified copy number, are molecularly ill-defined, and impossible to replicate in the quest to create derivatives with improved features. Thus, improving the ability to control expression levels in defined single copy transgenes would provide significant benefit to the transgenesis pipeline.

One transgenic approach which has been used widely to increase expression levels is the use of bipartite report systems including GAL4/UAS, Tet-On/*tetO* Tet-Off/*tetO*, QF/QUAS and lexA/*lexO* systems (Gossen and Bujard, 1992; Brand and Perrimon, 1993; Fashena et al., 2000; Potter et al., 2010; Yagi et al., 2010). These two component transgenic approaches allow for the amplification of weak promoters to permit higher expression levels than is feasible with direct promoter::reporter fusions. In addition, they are modular in design and thus can provide increased utility by being used in different combinations. Furthermore, the expression level of such transgenes can be tuned by altering the number of binding sites in the reporter promoter. Such systems have been reliably used in both invertebrate and vertebrate model systems. Notably, the Tet-On/Tet-Off system is widely used in mouse models (Schönig et al., 2010) and the GAL4/UAS system (and other bipartite systems as well) have been especially well refined in *Drosophila* and applied to many types of experimental approaches including genetic perturbation in conjunction with RNAi (Caygill and Brand, 2016; Riabinina and Potter, 2016).

Bipartite systems have been developed in *C. elegans* but have not been widely adopted. Although *C. elegans* would appear to be an ideal organism to use such reporters, initial attempts to port the GAL4/*UAS* systems to *C. elegans* were unsuccessful likely due to two factors 1) *S. cerevisiae* GAL4 functions poorly at low temperature (Wang et al., 2017), and 2) VP64 (4X derivative of VP16) is a very poor activator in *C. elegans* (Mao et al., 2019; Nonet, 2021). The QF/QUAS system was shown to function well in *C. elegans* including in single copy (Wei et al., 2012), but unfortunately the use of that system was not adopted by the research community. More recently, a derivative of the GAL4 system that uses *S. kudriavzevii* GAL4 (called cGAL4) was developed which works in multi-copy (Wang et al., 2017), but much less efficiently in single copy (Nonet, 2021). In contrast, another derivative that replaces VP64 with the QF activation domain works well in single copy (Nonet, 2020; Nonet, 2021). Additionally, derivatives of the Tet-On/*tetO* and Tet-Off/*tetO* system that also utilize QF as an activation domain were shown to function in multi-copy (Mao et al., 2019) and subsequently in single copy (Nonet, 2020). Furthermore, the lexA/*lexO* system has also been demonstrated to function in single copy (Nonet, 2020). Although these systems are functional, little work has been performed to assess the parameters that influence the functionality of these system i*n vivo*. What is the ideal number of binding sites for optimizing expression? What basal promoters function well with the enhancers? Is there tissue specificity to the expression levels driven by these systems?

Here we characterize some of parameters of *C. elegans* bipartite systems that influence expression. We examine how driver binding site number and sequence complexity influence expression for all four bipartite systems. We then focus primarily on the Tet-Off/*tetO* system to optimize additional parameters of the system. We examine how temperature influences expression and how expression profiles are influenced by tissue type. In addition, we address how binding site spacing influence expression. We test the influence of multiple basal promoters and 3’ UTRs on expression levels as well as the efficiency of transcriptional termination. Finally, we note that at least *tetO* reporters promote strong transcription in not only the forward direction, but also in the opposite orientation. Our analysis provides a knowledge base to aid in the rational design of reporters and an array of molecular components to assemble novel reporters. Furthermore, our work describes a variety of transgenic tools to facilitate testing the behavior of other drivers expressed under the control promoters that express in tissue types that have not been previously examined. It is hoped that these findings will promote the use of bipartite reporter systems in worms.

## Results

### Influence of *tetO* binding site number on expression

To optimize bipartite reporter systems, we initially concentrated on characterizing the influence of *tetO* binding site number on expression of a GFP reporter in touch receptor neurons (TRNs). The initial *tetO* fragment used to develop a single copy Tet-Off/*tetO* system consisted of seven copies of a slightly degenerate 36 bp sequence all containing the identical 19 bp *tetO* operator sequence (Nonet, 2020, **Fig. 1A**). We used this 7X *tetO* repeat as a building block to assemble 14X, 21X and 28X *tetO* promoters, and we truncated the 7X *tetO* fragment to create 1X, 2X, and 4X *tetO* modules. All *tetO* repeats were placed 5’ of a *mec-7* basal promoter (Nonet, 2020) driving GFP-C1 with a *tbb-2* 3’ UTR (**Fig. 1A**). The 2X, 4X, 7X, and 14X bipartite reporters were integrated on Chr II and crossed to a *mec-4p* Tet-Off driver transgene. At room temperature (RT, 22.5°C), expression of this reporter showed a concave down expression pattern with the highest expression obtained from a 4X reporter (**Fig. 1B**). Expression from the 14X promoter was virtually undetectable suggesting that large repeat numbers silence promoters (**Fig. 1B**). To further characterize the affects, we integrated all 7 reporters on Chr I and crossed them to the same Tet-Off driver. Quantification of expression (**Fig. 1C**) revealed that expression in TRNs was strongest with a 4X promoter at RT, reasonably robust from the 2X and 7X promoters, and weak from the 14X promoter paralleling the results from the Chr II transgenes. Expression was virtually undetectable from the 1X, 21X and 28X promoters. We then examined expression at other temperatures (**Fig. 1C**). A relatively similar expression profile was observed at 20° C and 25°C. However, at 15° C the 14X, 21X, and 28X promoters expressed much more comparably to the 4X promoter, though expression in general was lower than at higher temperatures. The expression profile at all temperatures was very similar in both PLM and ALM neurons (**Fig. S1**). These data indicate that there is a small range of binding sites that yields optimal expression at least in TRNs with reporters becoming increasingly silenced as binding site number increases above 14.

**Figure 1.**
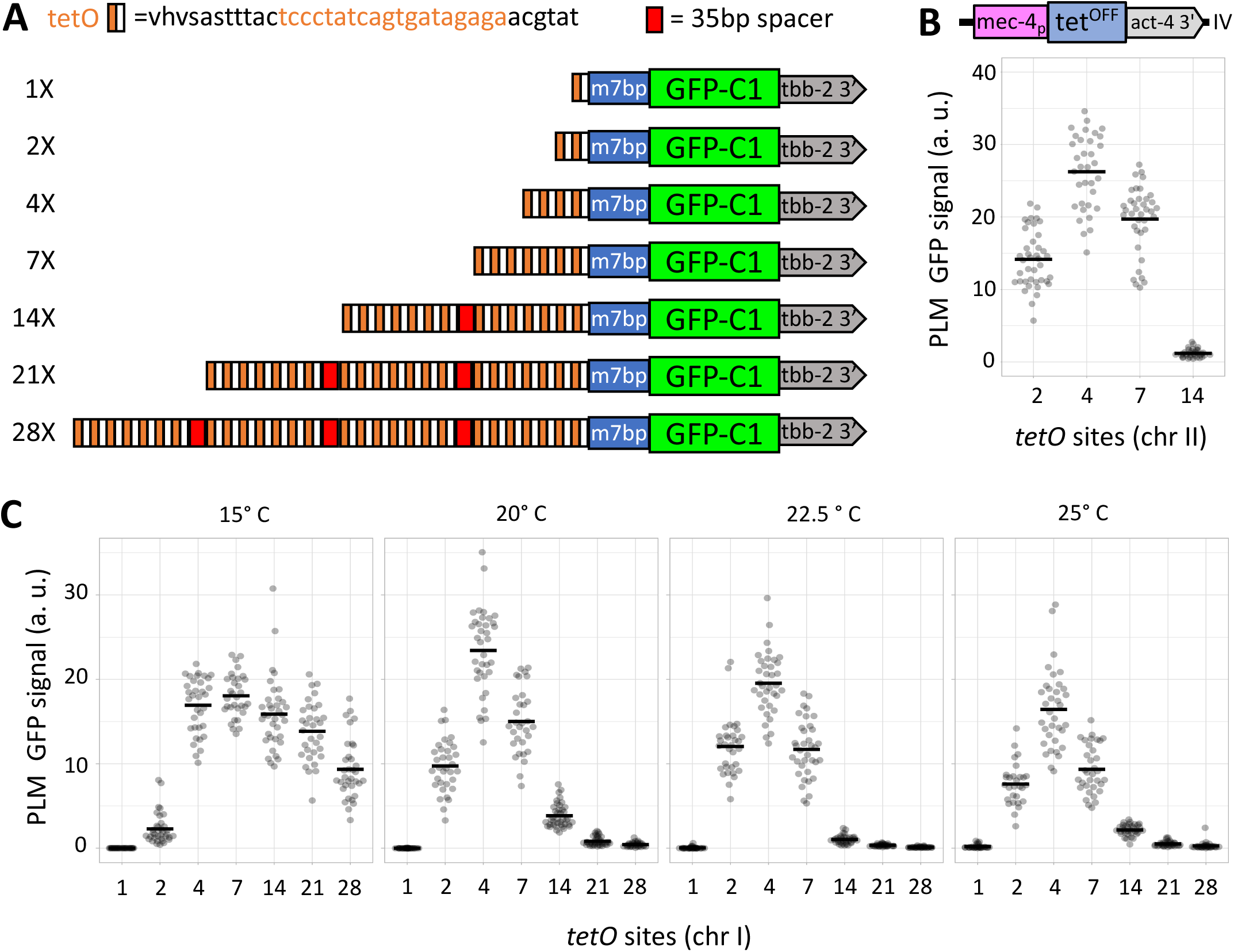
The influence of *tetO* copy number on expression. **A)** Schematic of the structure of single copy GFP reporters driven by promoters with varying number of *tetO* sites used to examine the effect of copy number on expression. The *tetO* binding sites and *mec-7* basal promoter (m7bp) are shown to scale, but GFP and the 3’ UTR are not. **B)** Quantification of expression levels in PLM neuron cell bodies from transgenic animals harboring reporter constructs with differing numbers of *tetO* sites driven by a *mec-4p* Tet-Off driver transgene integrated on Chr IV at 22.5° C (n=32-36). The structure of the driver transgene is shown above the graph. **C)** The influence of temperature on expression from a larger set of *tetO* reporters integrated on Chr I. All strains quantified are homozygous for both the driver and reporter (n=29-40). Genotypes of the strains used are listed in Table S1. Arbitrary units shown in B, C are defined identically and comparable.

To address whether the silencing effects observed with *tetO* binding site was a general property of bipartite systems in *C. elegans*, we created *UAS*, *QUAS* and *lexO* site repeats of varying size (**Fig. 2A**) and examined the expression of these reporters when a cognate driver was expressed in TRNs under the *mec-4* promoter. All reporters were integrated on Chr I at the same position as the *tetO* reporters. In all cases expression was maximal in the 8 to 10 binding site interval with lower expression at low and high binding site numbers (**Fig. 2B, S2A**). While *lexO* promoters were completely silenced at high repeat numbers similar to the *tetO* series, the QUAS and UAS promoters with high binding site numbers retained significant activity, though expression was reduced.

**Figure 2.**
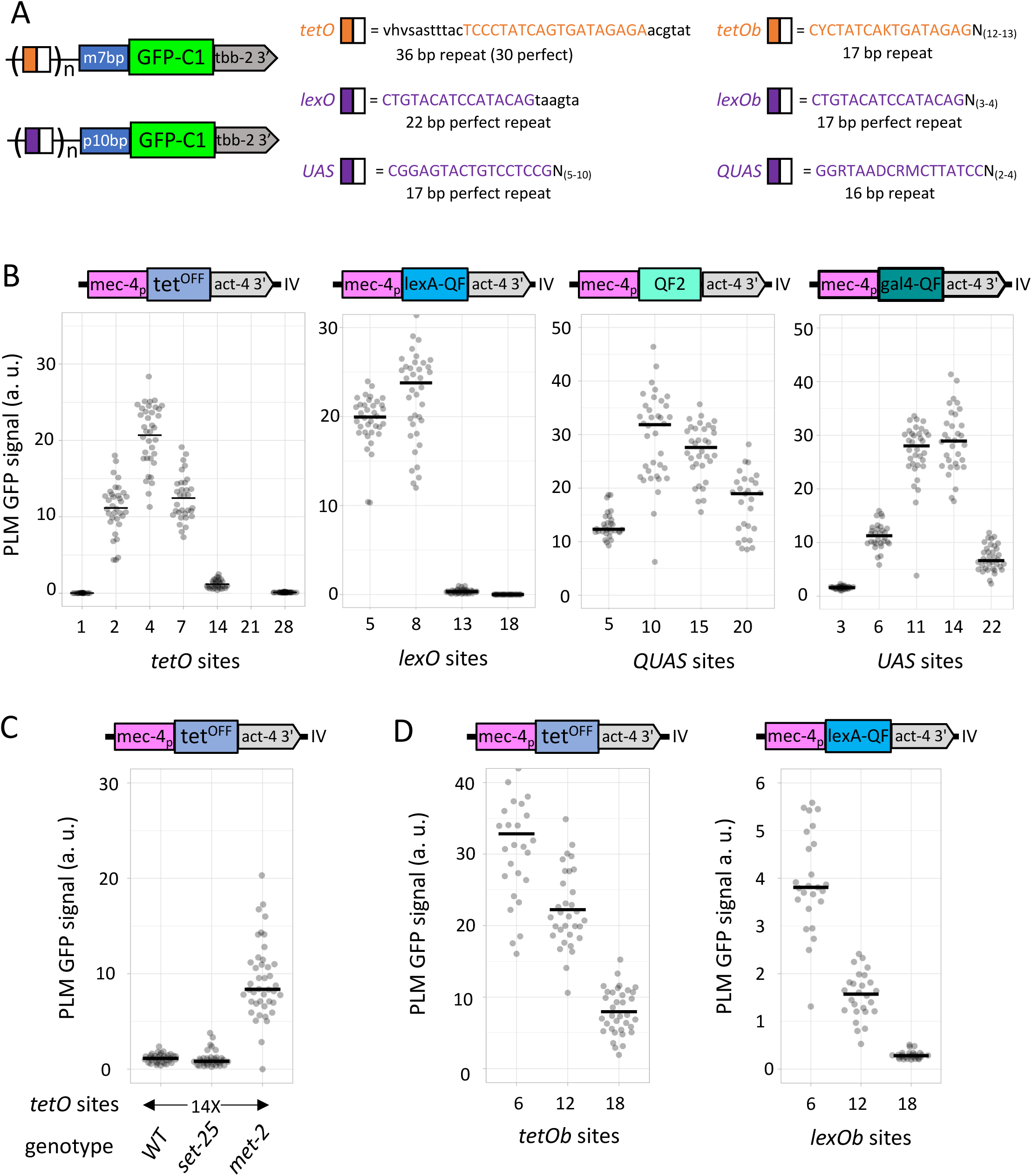
Influence of binding site complexity on expression. A) Schematic of the general structure of single copy GFP reporters used to examine the effect of repeat complexity on expression. Left) Schematics of the two basal promoter types used in the experiment. Right) Sequence of the six distinct binding sites examined. The sequence of *tetO* binding sites (UPPER case orange) placed upstream of the *mec-7* basal promoter (*m7bp*) and of the *UAS*, *QUAS* and *lexO* sites (UPPER case purple) placed upstream of the *pes-10* basal promoter (*p10bp*). Spacer sequences are shown in black lower case. **B)** Quantification of expression from reporters in PLM TRNs with varying number of binding sites driven by their cognate driver expressed under a *mec-4* promoter. Note that the less-repetitive binding sites support expression in higher copy number. No data was collected for the 21X *tetO* strain (n=27-40). **C)** Influence of chromatin modification defects on expression of *tetO* reporters. Quantification of expression from animals homozygous for a 14X *tetO* reporter, a *mec-4p* Tet-Off driver and for the chromatin modification mutations listed (n=36-42). **D)** Quantification of expression from reporters with varying number of minimally repetitive binding sites (shown in A) driven by their cognate Tet-Off driver expressed under a *mec-4* promoter (n=25-35). Arbitrary units shown in B, C and D are defined identically to Figure 1 and comparable. However, the units are not defined identically to those in Figures 3, 4, 5, and 6. Genotypes of the strains used are listed in Table S1.

### High *tetO* binding site promoters are de-silenced by lesioning chromatin modification machinery

One notable difference between the *tetO* and *lexO* binding sites versus the *QUAS* and *UAS* binding sites is the length of the perfect repeat unit. While *UAS* and *QUAS* sites consist of 17 bases or less and also contain a non-repeated component, the *tetO* and *lexO* repeats are larger (22 bases and 36 bases) and have little or no non-repetitive component (**Fig. 2A**). Silencing of transgenes in *C. elegans* that express in both somatic tissue and germline tissue has been studied in repetitive arrays and numerous factors which regulate silencing have been identified (Ahringer and Gasser, 2018). These factors include chromatin modification enzymes such as histone methyltransferases, chromatin structural elements, and components of small RNA processing pathways. We reasoned that the larger repeats found in the *tetO* and *lexO* promoters might be recognized by small RNA pathways and targeted for silencing when present in high copy. To test this hypothesis, a 14X *tetO* reporter and *mec-4p Tet-Off* driver were crossed into both *met-2* H3K9me1/2 histone methyltransferase and *set-25* H3K9me1/2/3 HMT mutant backgrounds, both of which have been reported to de-silence repetitive extrachromosomal arrays (Towbin et al., 2012). Disruption of *set-25*, but not of *met-2,* greatly increased the expression from the 14X *tetO* reporter indicating that chromatin structure is influencing expression from reporters with high *tetO* repeat number (**Fig. 2C, S2B**). To test if the silencing was the result of the length of the repetitive *tetO* binding site, we constructed synthetic 6X *tetO* DNA fragments with 17 bp binding sites separated by N_(12-13)_ (**Fig. 2A**). 6X, 12X and 18X reporter transgenes were created using 6X *tetO* modules and these were tested for expression by introducing the *mec-4p* Tet-Off driver. In contrast to original 14X 36 bp repeat reporter which is virtually completely silenced, a 18X 17bp repeat reporter was still expressed at similar levels to a 14X reporter in a *set-25* background and behaved more similarly to the QUAS and UAS reporters (**Fig 2D**). We also performed similar modifications to a *lexO r*eporter replacing the 6 bp repeat that separates *lexO* binding sites with a N_(3-4)_ spacer (**Fig 2A**). 6X, 12X and 18X reporters were created with this 6X *lexO* module. When crossed to a QF2 driver, we observed a similar pattern of de-repressed expression from the 12X and 18X lexO reporter (**Fig. 2D**). However, the maximal expression level from this set of promoters was greater than 10-fold weaker than the original *lexO* promoters suggesting that reducing the spacing between *lexO* binding sites from 22 bp to 19-20 bp was detrimental. Consistent with that interpretation, a 13X *lexO* reporter with 43 bp of non-repetitive sequence between *lexO* sites expressed much more robustly than the 12X reporter with 3 bp spacing between sites. Overall, these experiments indicated that highly repetitive binding site enhancer modules are more prone to silencing than less-repetitive ones.

### Effects of spacing of *tetO* binding sites on expression

To assess how spacing between individual *tetO* binding sites influences expression from reporters we first constructed a set of four different 4X *tetO* binding site modules separated by 3 bp, 25 bp, 43 bp and 93 bp (**Fig 3A**). To avoid the potential repressive influence of long repeats, we opted to create these DNA fragments by substituting the sequence of *tetO* sites in a ‘random’ 400 bp sequence which has the GC content and motif bias of *C. elegans* promoter sequences (see supplemental methods for details). The four binding site modules were inserted upstream of a *mec-7* basal promoter and GFP-C1 reporter and tested for expression by crossing to both a TRN and an intestinal driver. In TRNs, increasing spacing from 3 to 25 or 43 bp modestly increased expression but extending the spacing to 93 bp dramatically reduced expression (**Fig 3A**). Similarly, in intestinal tissue, expression of reporters with 93 bp spacing was very weak, but by contrast with TRNs, 3 and 25 bp spacing was over two-fold less effective than 43 bp spacing.

**Figure 3.**
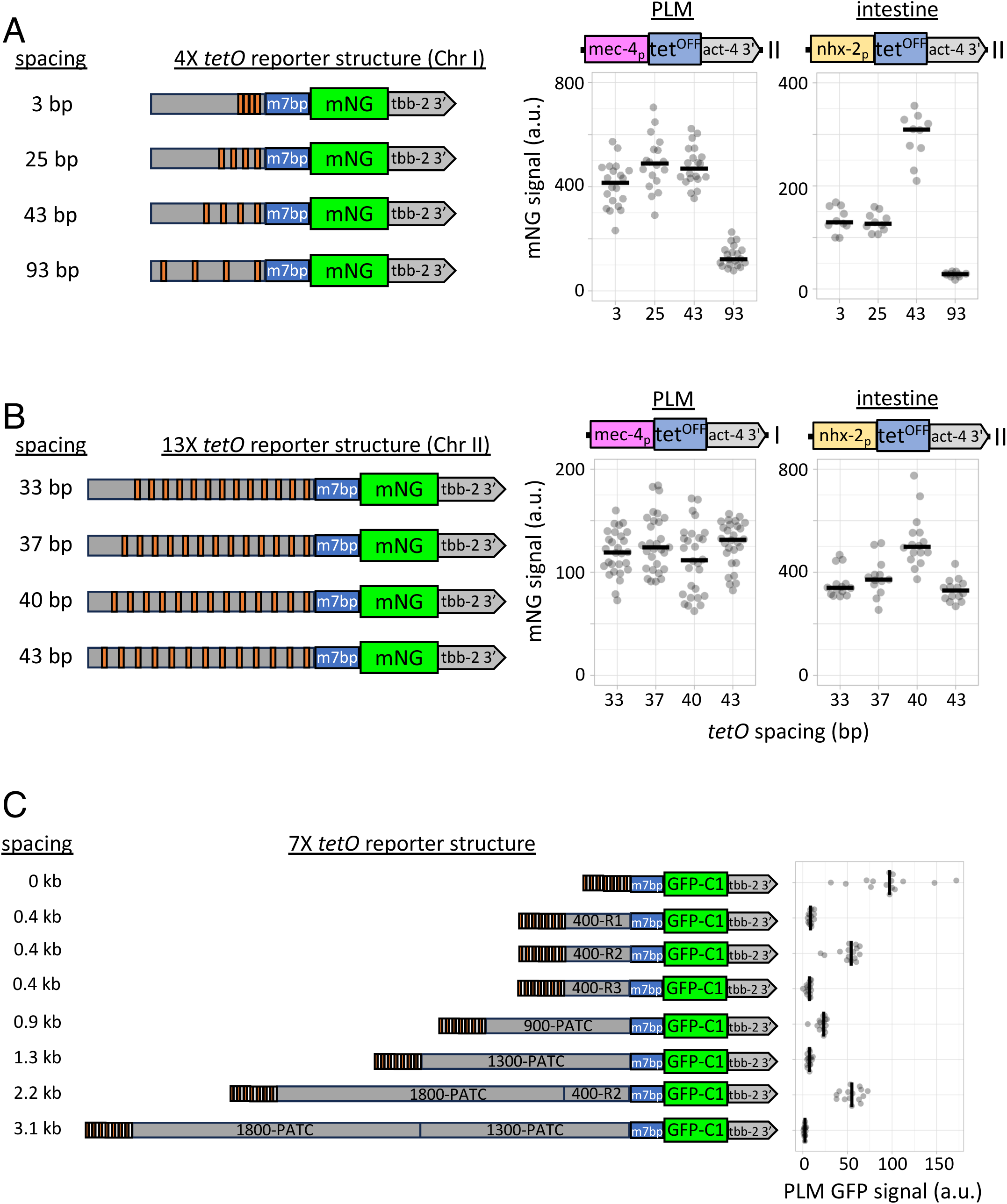
Influence of *tetO* site spacing on expression. **A)** Left) Schematic of the structure of single copy mNG reporters carrying four *tetO* sites positioned with varying spacing within the same 400 bp randomly generated sequence with a GC content and sequence composition similar to *C. elegans* promoters. Right) Quantification of expression levels in both PLM neurons and the intestine as a function of spacing. Schematic of drivers used in strain construction are shown above the graph. All animals are homozygous for both the driver and the reporter (*mec-4p*: n=18-20, *nhx-2p*: n=10). **B)** Left) Schematic of the structure of single copy mNG reporters carrying thirteen *tetO* sites positioned with varying spacing within the same 800 bp randomly generated sequence with a GC content and sequence composition similar to *C. elegans* promoters. Right) Quantification of expression levels in both PLM neurons and the intestine as a function of spacing. Schematic of drivers used in strain construction are shown above the graph. *mec-4p* driver/reporter animals are homozygous for both the driver and the reporter (n=28-29) and *nhx-2p* driver/reporter animals are *trans* heterozygotes (n=13-16). **C)** Left) Schematic of the structure of single copy mNG reporters carrying seven *tetO* sites positioned with varying distance from a *mec-7* basal promoter (*m7bp*). The spacers consist of PATC rich sequences from the *C. elegans* genome and 400 bp random sequences as described in A. See supplemental methods for details. Right) Quantification of expression levels in PLM neurons as a function of spacing between the basal promoter and the *tetO* enhancer. The *tetO* binding sites and *mec-7* basal promoter (*m7bp*) and spacers are shown to scale, but GFP and the 3’ UTR are not. Genotypes of the strains used are listed in Table S1.

Double stranded DNA has a helical repeat of 10 bp. To assess if the position of *tetO* binding sites on the DNA relative to the helical repeat influences expression, we created 13X *tetO* binding site modules with spacing of 33 bp, 37 bp, 40 bp and 43 bp between *tetO* sites (**Fig. 3B**) with the logic that one of these four would likely closely approximate an optimum that, regardless of what specific spacing, was ideal. The modules were introduced into reporter constructs as outlined above and tested by crossing to TRN and intestinal drivers. Although we observed some differences in expression between the different modules, no consistent pattern was observed in the cell types we examined and the differences between the weakest and strongest was modest (**Fig. 3B**). While the position of *tetO* binding sites relative to the helical repeat had minimal effects, we conclude that spacing has strong influence on expression. Specifically, it was dramatically reduced by extended spacing (∼100 bp) but less strongly influenced by changes in spacing in the 25 to 50 bp range. Very close spacing is also undesirable because it does not afford the ability to disrupt the repeats and it greatly complicated the *in vitro* synthesis and PCR amplification of the binding site modules.

*C. elegans* does not contain homologs of the classic insulator proteins and thus isolation of individual promoters from neighboring genes has been proposed to occur primarily due to spacing (Heger et al., 2009). Whole genome level analysis of expression characteristics of neighboring genes supports this model (Quintero-Cadena and Sternberg, 2016). Thus, we examined how spacing between the basal promoter and the *tetO* binding site module influences expression. We used a combination of random sequences with sequence complexity similar to *C. elegans* promoter regions and <u>p</u>eriodic <u>A</u>n/<u>T</u>n <u>C</u>lusters (PATC) rich regions (Fire et al., 2006) that constitute 6.1% of the *C. elegans* genome as spacers between a *mec-7* basal promoter and a 7X *tetO* module (**Fig 3C**). Placing random 400 bp sequences with sequence complexity and GC content similar to *C. elegans* promoters reduced the efficacy of the *tetO* binding site enhancer to drive transcription to variable extent. 900 and 1300 bp PATC-rich regions had a similar effect on expression as the smaller random promoter sequences. Increasing the spacing to 3.1 kb only reduced but did not eliminate expression suggesting that spacing that is much greater than the median spacing between divergent promoters (∼950 bp) in *C. elegans* is not always sufficient to isolate a promoter from a regulatory element. Indeed, a 2.2 kb spacer between the basal promoter and the tetO 7X enhancer consisting of a 1800 bp PATC region the 400-R2 bp random sequence was virtually as effective as the 400-R2 sequence alone suggesting that some sequences are capable of fostering enhancer promoter interactions even at considerable distances.

### *tetO* site number expression profile is influenced by tissue type

To investigate how tissue type influences the Tet-Off/*tetO* bipartite reporter system, we created other Tet-Off driver constructs that express in neurons, body wall muscle, pharyngeal muscle, coelomocytes, the intestine and epidermal tissue using well characterized promoters (**Fig. 4A**). The driver transgenes were crossed to the 2X, 4X, 7X, 14X, 21X, and 28X *tetO* reporter lines and the expression level in each tissue was quantified at RT (**Fig. 4B and S4**). Expression in the epidermis, pharyngeal and body wall muscle, and the coelomocytes increased with repeat number until 7X then largely plateaued from 7X to 28X. By contrast, pan neuronal and intestinal expression increased until 14X, but then declined substantially at 21X and 28X. However, all the tissue specific expression profiles were much less dramatically silenced at high repeat numbers than the TRN profile. The promoters used for these tissue specific studies are all quite strong. Thus, we tested additional promoters to test the influence of promoter strength on expression. We tested a synthetic *mec-4* promoter which increases expression 2.5-fold over the native *mec-4* promoter (Dour and Nonet, 2021). Expression using the synthetic reporter mimicked the pan-neuronal expression series, declining substantially at 21X and 28X while retaining some expression (**Fig. 4B**). We also examined the *nhx-2* promoter [6-fold weaker than *vha-6p* as assayed by RNAseq (Cao et al., 2017)] and this promoter exhibited a very similar profile to the *vha-6* promoter (**Fig. S4**). Finally, we tested the *phat-5* promoter which is expressed exceedingly poorly in single copy (Nonet, 2020). The *phat-5* driver expressed very weakly with 2X and 4X t*etO* promoters and was exceedingly variable with 7X-28X *tetO* promoters (**Fig. S4**). Our data indicate that both promoter strength and tissue type are likely to influence the expression levels obtained with any specific *tetO* reporter. In practical terms, our data support the strategy of using a promoter containing seven *tetO* binding sites which is likely to express at consistently high levels in most tissues and probably represents the most suitable option for creating reporters to be used in multiple different cell types. However, especially in the case of weak promoters, if optimal expression is desired in another tissue, then performing a series of test crosses with reporters containing differing numbers of binding sites could provide substantially higher expression levels than a 7X promoter.

**Figure 4.**
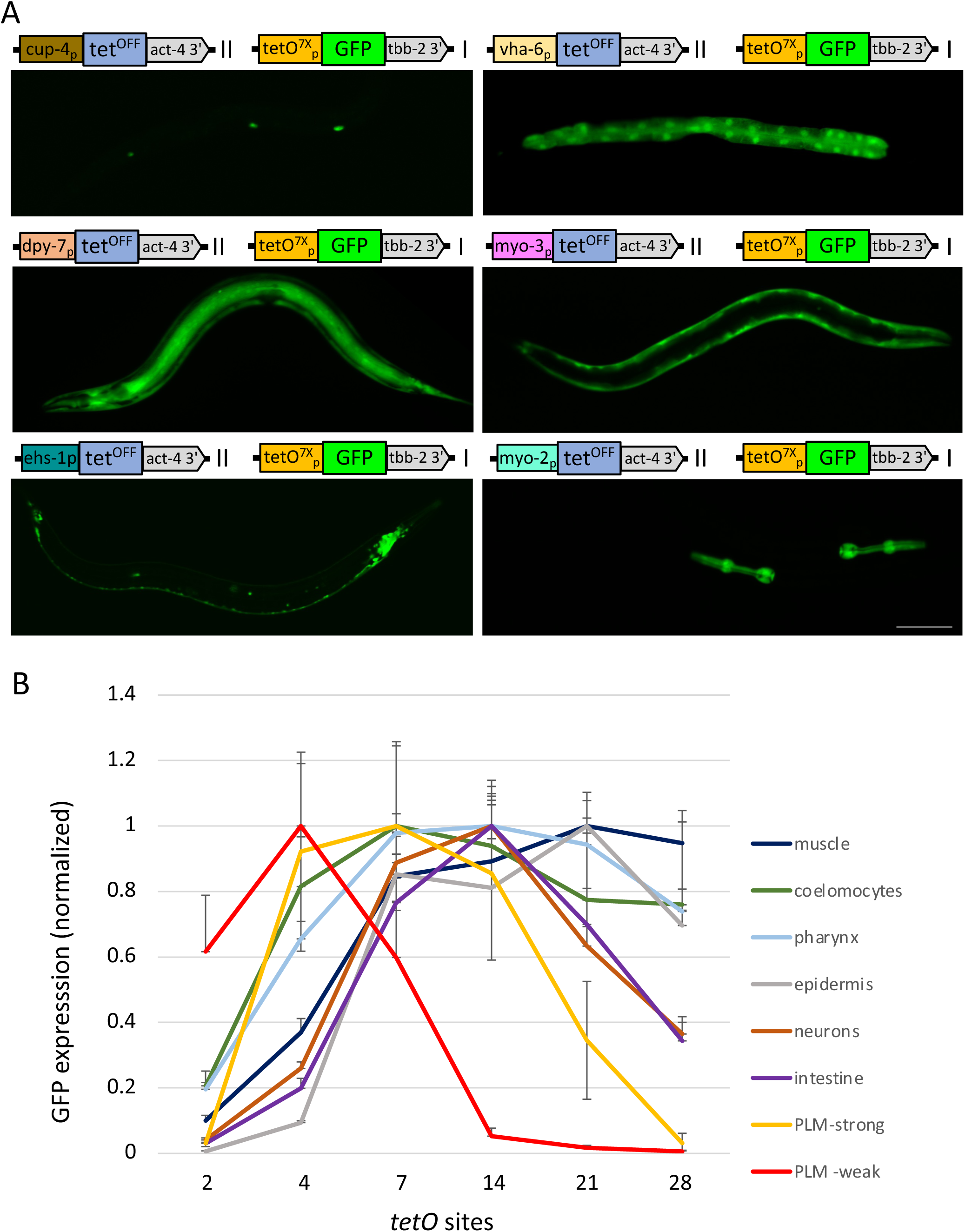
Tissue specificity of *tetO* expression profiles. **A)** Schematic of the transgenes present in animals harboring both tissue-specific Tet-Off drivers and 7X *tetO* reporters and representative images illustrating the expression pattern in L4 animals homozygous for both the driver and the reporter. **B)** Normalized expression levels from reporters with different *tetO* copy number driven by distinct tissue-specific Tet-Off drivers. Error bars represent S.D. See Figure S4 for plots of the individual tissue data. Genotypes of the strains used are listed in Table S1. Scale Bar: 100 µm.

### Influence of basal promoter on expression levels and specificity

We previously tested several basal promoters for use in conjunction with repetitive *tetO* and *UAS* promoters (Nonet, 2020). To extend our understanding of the influence of basal promoters on expression, we tested additional basal promoters in the context of a *7X tetO* enhancer driving GFP-C1. We chose three promoters that have well defined transcriptional start sites (Gu et al., 2012; Chen et al., 2013; Kruesi et al., 2013; Saito et al., 2013) and either express strongly in single copy transgenes (*lys-8p*) or transcribe genes which express highly abundant actin and tubulin subunits (*act-4* and *tbb-2*). In addition, we tested the *pals-5* promoter which is dramatically induced in the gut during pathogen infection (Bakowski et al., 2014; Chen et al., 2017). Bioinformatic analysis of promoters has not revealed many commonalities shared among their sequences (Grishkevich et al., 2011; Reinke et al., 2013; Saito et al., 2013). Numerous motifs have been identified but are each only present in less that 5% of promoters. Thus, we relied on earlier studies which typically found that the sequences 100 to 200 bp upstream of the ATG functioned as’ basal’ promoters in that they did not drive expression in the pattern of the entire regulatory region, but were capable of expression at high level when placed upstream of a distinct regulatory region (Okkema et al., 1993; Duggan et al., 1998; Thatcher et al., 2001; Hong et al., 2004; Arribere et al., 2014). After testing an initial set of promoter fragments, we either extended or reduced the size of the promoter with the aim of optimizing expression levels in the presence of a driver and minimizing background expression in the absence of the driver. All the basal promoters we examined showed some level of expression in absence of a driver, though it was, in most cases, a relatively low level and restricted to a small subset of cells (**Fig. 5A and S5**). The level of expression in the presence of tissue specific drivers varied over 10-fold depending on the basal promoter. In addition, the basal promoters exhibited some tissue specificity. For example, the *mec-7* basal promoter functioned much better in TRNs than in intestinal tissue, while the *tbb-2* basal promoter showed the opposite preference. Our work with transgenes has previously shown that promoters of adjacent transcriptional units can exhibit significant crosstalk (Nonet, 2023). Thus, we also examined the basal promoter’s susceptibility to influence by neighboring regulatory elements. In mechanosensory neurons, the presence of an upstream *myo-2p::Scarlet* and *rps-0::Hyg^R^*cassette reduced the expression of the *tetO* basal promoter expression in 3 independent cases. By contrast, in intestinal tissue, there was no influence of the adjacent transcription units on expression. In summary, several additional basal promoters with strong expression profiles were characterized and these promoters behave distinctly based on tissue type and local genomic environment.

**Figure 5.**
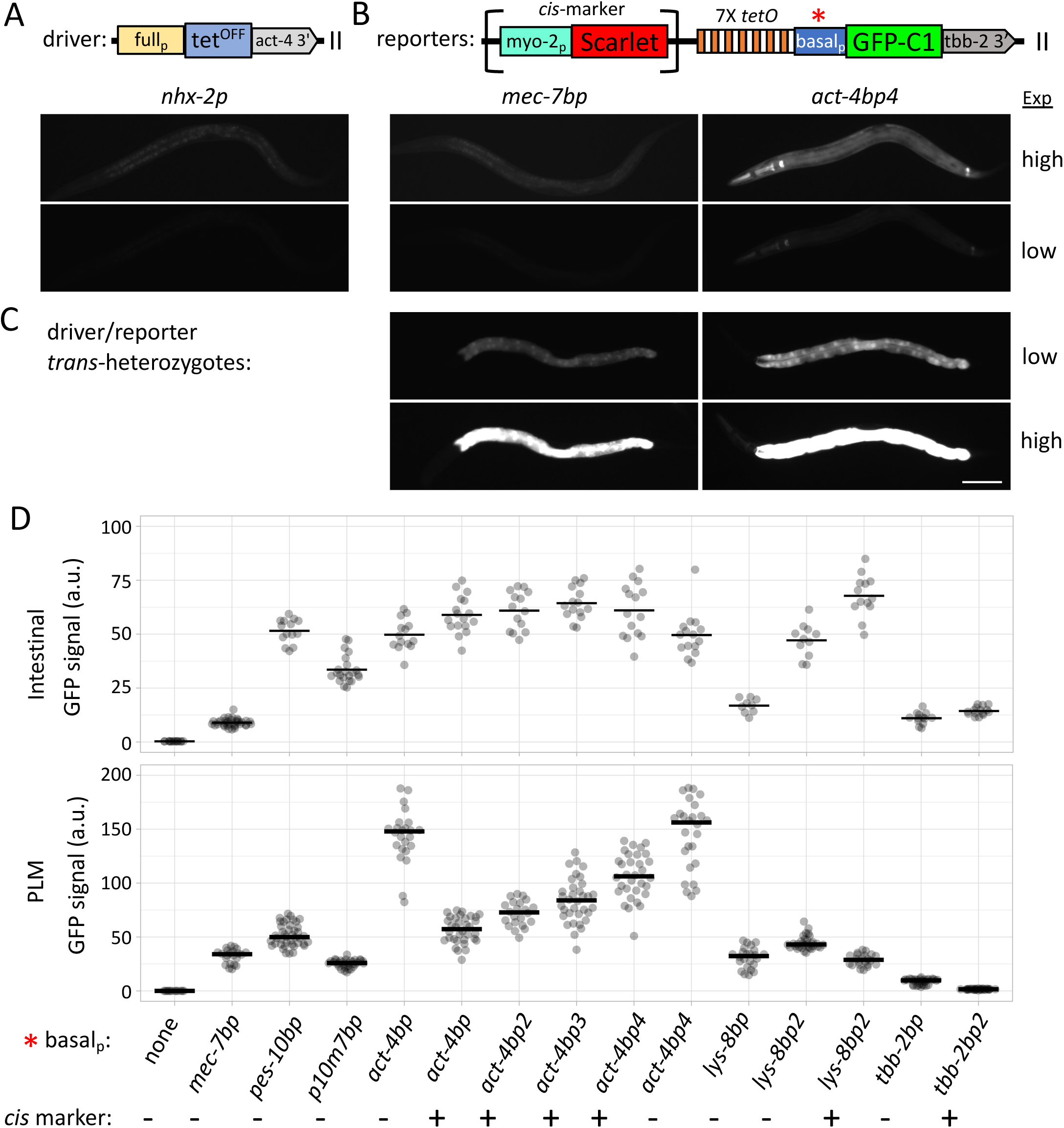
Functionality of basal promoters in *tetO*/Tet-Off bipartite expression. **A)** Schematic of Tet-Off driver transgenic insertions used to examine basal promoter activity. Below the schematic is an image of an L4 animal homozygous for an *nhx-2p* intestinal driver insertion. The image shows the level of auto fluorescent signal present in *C. elegans* animals not containing a FP. **B)** Schematic of the general structure of the *tetO* reporter transgenic insertion*s* used to examine promoter activity of various basal promoters. The red asterisk shows the position of the basal promoter. Some insertions made using rRMCE mediated integration contain a *cis*-linked marker shown surrounded by brackets. Below the schematic are short and long exposures of L4 animals taken in a GFP channel showing the level of background associated with the basal promoter with the lowest background (*mec-7bp*) and the highest background (*act-4bp4*) of all basal promoters analyzed. Similar images for the other basal promoters are shown in Table S5. **C)** Images of L4 *trans* heterozygous animals carrying one copy of the *nhx-2p* Tet-Off driver and one copy of a *7X tetO* GFP reporter harboring either the *mec-7bp* or *act-4bp4* basal promoter. The exposures in C) are 10-fold shorter than those in B). **D)** Quantification of GFP signal in *trans* heterozygote L4 animals containing one copy of either an intestinal-specific *nhx-2p* driver (top) or a TRN-specific *mec-4p* driver and one copy of a *7X tetO* GFP reporter of structure schematized in B) harboring a basal promoter as indicated. In some cases, both reporter strains with the *cis*-marker present and the *cis*-marker excised were analyzed (*nhx-2p*:n=9-35, *mec-4p*: n=22-42). Genotypes of the strains used are listed in Table S1.

### Random sequences function as basal promoters

Since bioinformatic studies found minimal conserved motifs among *C. elegans* basal promoters, we explored the possibility that transcription initiation can occur relatively non-specifically independent of any specific sequence motif(s). We generated random 200 bp sequences with GC content and composition typical of *C. elegans* promoters using a random sequence generator (Thomas-Chollier et al., 2011). We filtered the set of random sequences for those without an ATG on the top strand to eliminate the possibility that an ATG upstream and out of frame of the GFP reporter open reading frame would confound our reporter assay. Three such sequences were tested as basal promoters (**Fig. 6A**) and each functioned to a modest extent, expressing at ∼ two thirds the efficiency of a comparable transgene containing the *mec-7* basal promoter. A 400 bp random sequence with similar sequence feature which was not filtered for ATGs performed less effectively, but still promoted some transcription. These data suggest that either transcription initiation does not require very strict DNA sequence motifs or that initiation is occurring from elsewhere, presumably within the *tetO* sequence region.

**Figure 6.**
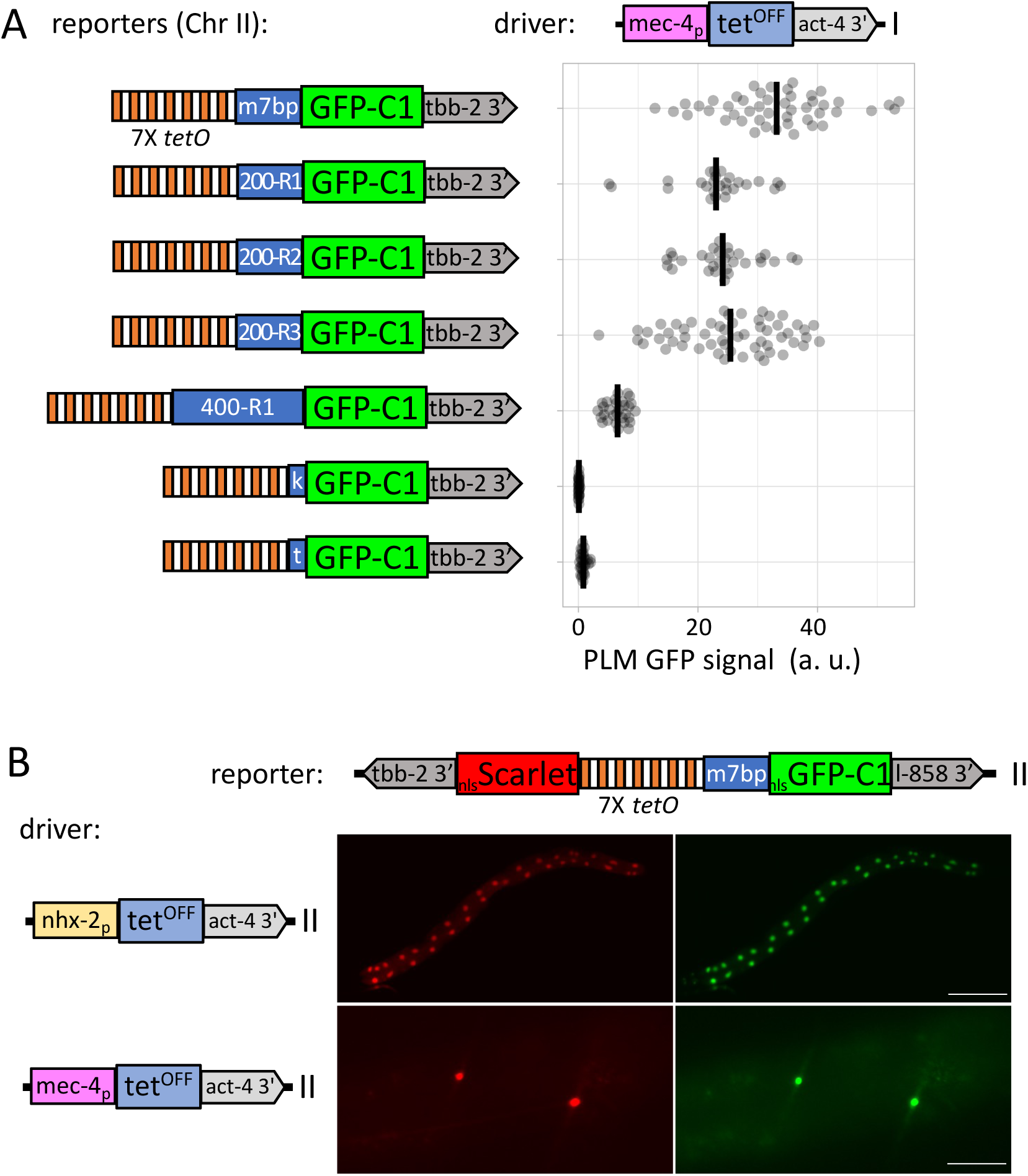
Additional properties of basal promoters. **A)** Left) A schematic of the structure of insertions with random 200 bp (rand1-rand3), 400 bp (rand4) or small sequence motifs in place of a traditional basal promoter. The motif (k) consists of a *C. elegans* consensus Kozak sequence (AAAA) directly upstream of the ATG. The small motif ‘t’ consists of an SL1 trans-splice signal and a Kozak (TTTCAGCAAAA) directly upstream of the ATG. Right) Quantification of expression in PLM from transgenes homozygous for both the reporters and a *mec-4p* driver schematized above the graph (n=22-56). **B)** Schematic of drivers and reporters used to assess if *7X tetO* based promoters drive bidirectional transcription. No basal promoter was introduced between the *7X tetO* repeat and the Scarlet reporter. Images of L4 *trans* heterozygotes carrying both a driver and the bidirectional reporter. The red and green channel images were taken with the same exposure time. Scale bars 5A: 100 µm. 5F top: 100 µm, bottom: 25 µm. Genotypes of the strains used are listed in Table S1.

Analysis of transcriptional initiation at a genome wide level has revealed that initiation is also often seen from enhancer regions in many organisms including *C. elegans* (Chen et al., 2013; Li et al., 2016; Akay et al., 2019). To assess if initiation might be occurring from within the *tetO* enhancer region, we constructed transgenes lacking a basal promoter. The sequences between the 7X *tetO* module and the ATG were limited to a 4 bp consensus Kozak signal or an 11 bp sequence including both an SL1 trans-splice signal and a closely spaced Kozak sequence. Expression from both of these transgenes lacking a basal promoter was minimal (**Fig. 6A**) suggesting that little transcription initiation is occurring from within the *tetO* binding site region.

### *tetO* promoters drive bidirectional transcription

In many systems including both invertebrates and vertebrates, transcription initiation is not exceedingly strand specific (Bagchi and Lyer, 2016). In addition to the expected forward transcription, significant transcriptional initiation occurring in the opposite direction at many promoters is also observed. This type of transcription is relatively common in *C. elegans* based on genomic analysis mapping of transcriptional start sites (TTSs) with 39% of transcriptional initial complexes being associated with reverse strand initiation with 200 bp upstream of the forward initiation site (Chen et al., 2013). We asked if a *7X tetO* promoter was functioning bidirectionally by inserting a Scarlet reporter in the opposite direction upstream of the *tetO* sequences. We did not include a second basal promoter in the construct. Robust transcription of Scarlet was observed both in mechanosensory neurons and the intestine when crossed to driver lines specific for these tissues (**Fig. 6B**). Transcription of Scarlet was easily detected on the dissecting scope and appeared roughly comparable to GFP expression. Exact quantification of the relative level of initiation in both directions is difficult to perform using fluorescent assays since different fluorophores are being expressed and different 3’ UTR are controlling transcript stability. Regardless of the exact relative level of transcription, the propensity for reverse direction transcription is clearly an event that must be considered in the design of transgenes.

### Influence of 3’ UTR on expression

Prior studies have documented the strong influence that 3’ UTRs can have on expression from transgenes (Merritt et al., 2008). However, only a few sequences have been routinely used in transgenic constructs including the *let-858, unc-54, tbb-2, and glh-2* 3’ UTR. Ideally, a 3’ UTR used in a transgene should permit high expression in all tissues, but not act in any capacity as a direct stimulator of transcription. In addition, a 3’ UTR should terminate transcription efficiently. In analyzing expression from bipartite transgenics, we noticed that many transgenic animals harboring the *tbb-2* 3’ UTR expressed in the pharynx suggesting that it might have enhancer activity. Thus, we tested both commonly used and additional 3’ UTRs for enhancer activity by placing them upstream of a basal promoter driving GFP and the *let-858* 3’ UTR. In this assay the *tbb-2* 3’ UTR expressed in pharyngeal muscle (**Fig. 7A)**. However, other 3’ UTRs including act*-4, glh-2, rps-0, rps-2, rps-9, ubl-1*, and *unc-54* exhibited no transcriptional activation properties (**Fig. 7A, 7S)**. The *let-858* sequence used as the 3’ UTR for the reporter itself exhibited a low level of expression in several cells surrounding the pharynx, which were not specifically identified, but are likely either of neuronal or glial origin (**Fig. 7A**). The *eef-2* 3’ UTR also exhibited a pattern of expression that was very similar to let-858 but included more cells (**Fig. 7A).** Thus, a significant fraction of 3’ UTRs exhibit some transcriptional enhancer activity including several commonly used in transgenic constructs.

**Figure 7.**
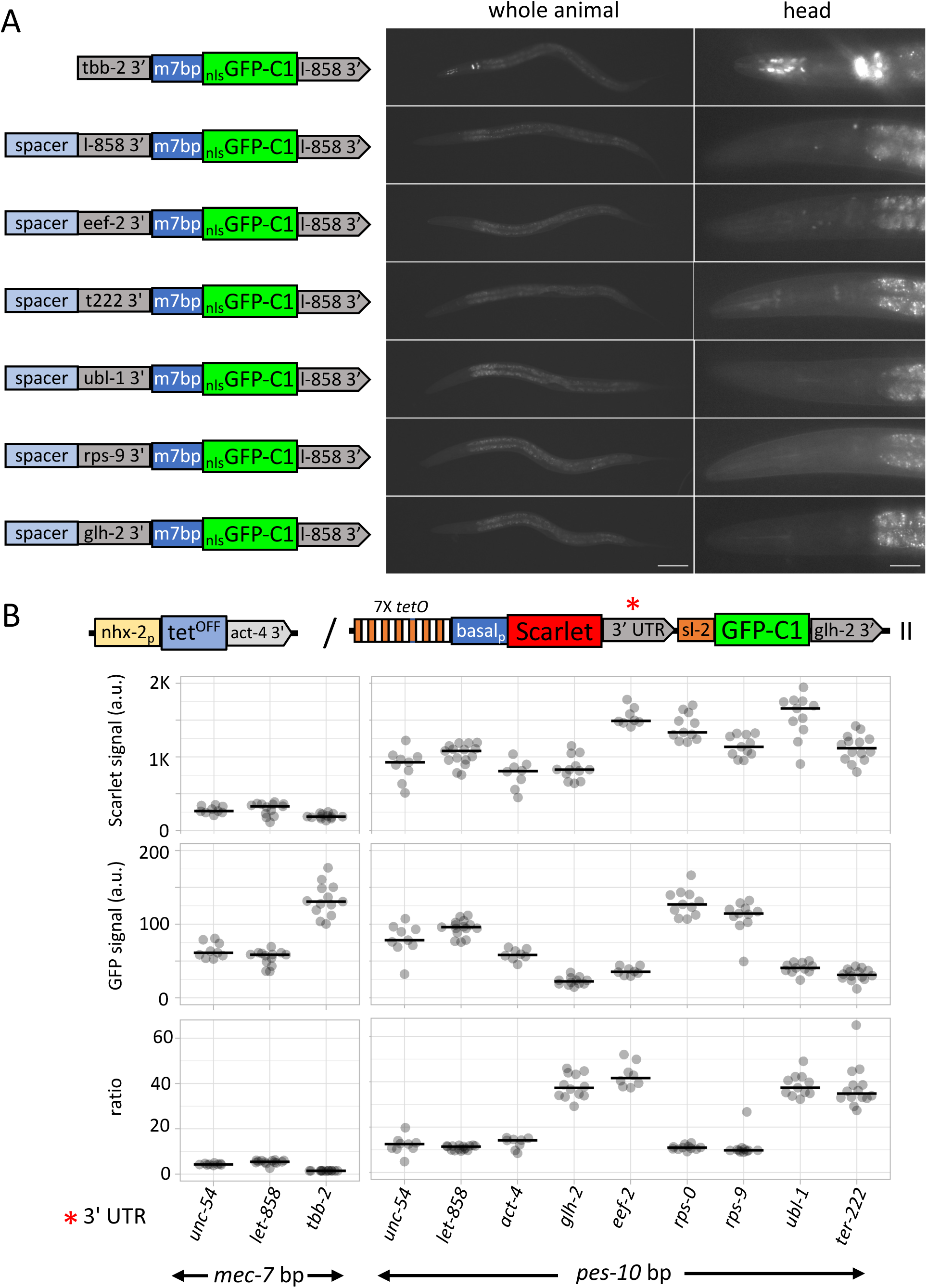
Influence of 3’ UTRs on expression from bipartite promoters. **A)** Left) Schematic of transgenic insertions used to test for promoter/enhancer function of commonly used 3’ UTRs. Right) Images of GFP fluorescence in animals homozygous for constructs illustrated on the left. Images of the whole body (left) and head region (right) of L4 animals. The genomic region used for each 3’ UTR are shown in Figure S7A. Images for additional 3’ UTRs are shown in Figure S7C. Scale bars, left column:100 µm, right column: 25 µm. **B)** Top) Schematic illustrating the structure of the Tet-Off driver and *7X tetO* reporters used to quantify the influence of 3 ‘ UTRs on both expression levels and the efficiency of transcriptional termination. A red asterisk denotes the position of the 3’ UTR in the reporter construct. Bottom) Quantification of Scarlet and GFP fluorescence in animals heterozygous for an *nhx-2p* intestinal driver and a reporter with varying 3’ UTRs (red asterisk) and basal promoters as indicated on the X-axis (n=9-15). The ratio of the Scarlet to GFP signal is also plotted. The ratio is not an absolute measure of the ratio of Scarlet to GFP molecules since the relative efficiency of fluorophore excitation and emission detection for the two channels was not determined. However, relative ratios should reflect the relative termination efficiency of different 3’ UTRs. Genotypes of the strains used are listed in Table S1.

To quantify the ability of 3’ UTRs to influence expression levels we created reporter transgenes containing a *7X tetO* module and a basal promoter driving Scarlet with distinct 3’ UTRs and crossed these to a *nhx-2p* intestinal driver. Expression levels varied over 5-fold depending on which basal promoter and 3’ UTR was utilized (**Fig. 7B)**. The expression levels generally paralleled that observed in similar studies in touch receptor neurons (Dour and Nonet, 2021).

One additional property of 3’ UTR sequences, including sequences downstream of the poly adenylation site, is their ability promote transcriptional termination. This process in *C. elegans* has been characterized in several studies and shown to occur using at least two distinct mechanisms (Haenni et al., 2009; Miki et al., 2017). Transcriptional termination is more complicated in *C. elegans* than many other metazoans because operons are common. In such multi-gene operons, cleavage and polyadenylation of the primary transcript occurs without RNA polymerase termination and the extending transcript is trans-spliced using a distinct leader from that used on non-operon genes (Graber et al., 2007). To test the efficiency of termination we used an assay similar to that developed by Miki et al. (2017) where the operonic *gpd-2/gpd-3* trans-splice leader sequence and a reporter are placed downstream of the 3’ UTR region. If the 3’ UTR region terminates transcription inefficiently then trans-splicing via the *gpd-2/gpd-3* signal yields transcription of the downstream reporter. The ratio of the upstream Scarlet reporter level to the downstream mNG reporter can be used to compare the relative termination efficiencies of distinct 3’ UTR regions. Our data show that transcription initiated under control of the strong *pes-10* basal promoter were of two classes. *glh-2*, *eef-2, ubl-1* and a synthetic triple termination consisting of the *tbb-2*, *rpl-2* and *rps-2* 3’ UTR in tandem all displayed a high ratio (**Fig. 7B**). In contrast, the more commonly used *unc-54*, *let-858* and *act-4* 3’ UTRs as well as the *rps-0* and *rps-9* 3’ UTRs show an approximately four-fold lower ratio suggesting these terminate less efficiently (**Fig. 7B**). It is expected that *rps-0* and *unc-54* would have inefficient termination as *rps-0* is the first gene of an operon and *aex-5* transcription initiates within the *unc-54* 3’ UTR region though it is not defined as an operon in Wormbase (Davis et al., 2022). However, it was unexpected that *rps-9, let-858* and *act-4* would terminate less efficiently. Our data suggest that *ubl-1* is likely a suitable 3’ UTR to use when aiming for high expression of transgenes.

## Discussion

Bipartite reporter systems are versatile tools for creating cell specific genetically encoded markers for dissecting molecular mechanisms. They provide the ability to amplify the signal from weak promoters and the efficiency of combinatorial systems. However, they are also complicated to develop as multiple parts of the system are changeable, including the number of activator binding sites, the basal promoter, and the 3’ UTR. Here we have performed basic characterization of several bipartite reporters to develop a framework for the rational development of reporter transgenes. In addition, we describe numerous molecular and genetic tools to simplify the development and characterization of additional bipartite reporters.

### Prior optimization

Several different groups have performed bipartite system optimization experiments by altering the number of driver binding sites in the reporter promoter. Those experiments have generally concluded that increasing binding sites eventually reduces expression, though the repeat number at which this occurs varies greatly by model and the bipartite system used (Agha-Mohammadi et al., 2004; Distel et al., 2009; Pfeiffer et al., 2010). In *C. elegans*, 15X UAS was determined to be more active in multicopy arrays than 5X, 10X or 20X (Wang et al., 2017).

Our results show a very similar pattern, with large repeat numbers reducing expression. Interestingly, in single copy transgenes using a Δmec-7 or a Δpes-10 basal promoter, the optimal repeat number is different for distinct bipartite systems being lower for lexA/lexO and tetOFF/tetO than for QUAS/QF2 and GAL4/UAS. Furthermore, for the *tetO* system the optimal repeat number is influenced by the expression level of the driver as well as the tissue type. The reporter lines with differing numbers of driver binding sites developed herein will provide a simple genetics-based approach to quickly determine the optimal repeat number for novel drivers expressing in either different tissues or using other promoters.

In addition to binding site number, binding spacing has also been demonstrated to greatly influence expression. In mammalian tissue culture, expression from 2X *tetO* promoters was much more robust with a half of a turn spacing between binding sites, placing TetR/DNA complexes on opposite sides of the helix, compared to sites on the same side (Agha-Mohammadi et al., 2004). We tested how both spacing and positioning along the helix influences expression levels. By contrast with Agha-Mohammadi et al. (2004), we found that expression varied as spacing between repeats in relation to helix turns varied, but this effect was not consistent between the two tissues examined. Several things could account for this. Since 2X *tetO* promoters express poorly in worms, we opted to examine 4X and 13X promoters. In these multi-binding site promoters, DNA looping between distally positioned sites may counteract the steric effects that are postulated to influence the ability of bound tetR-QF dimers to activate transcription initiation. Alternatively, the larger size of the QF activation domain used in our studies versus the smaller VP16 domain used in the Agha-Mohammadi led studies may provide more flexibility in the positioning of binding sites. Regardless of what mechanism underlies the differences, our data cannot be converted into clear recommendations for how to space *tetO* binding sites. We did not test how the spacing of *lexO*, *QUAS* or *UAS* binding sites influence expression though we have documented that 13X *lexO* and 13X *QUAS* binding sites separated by 43 bp are functional (not shown).

Our original impetus for examining repeat spacing was to ameliorate the silencing effects we observed with tetOFF/*tetO* reporters in TRNs. However, an additional benefit of spacing the *tetO* sites further apart is the potential to introduce other sequences between the sites. This could be valuable for fine-tuning the strength of reporters, such as in cases where genetically encoded organelle markers spill into other compartments when highly expressed. Introducing efficient sgRNA sites between binding sites would enable one to delete regions of the *tetO* site cluster to reduce expression. With recent advances in genetically encoding Cas9 and sgRNAs (Schwartz et al., 2021), such ‘tuning’ of reporters might be feasible through simple crosses to transgenic lines expressing both cas9 and sgRNA(s).

### Transcriptional Drivers

Although this work examined the influence of many aspects of reporter structure on expression, it largely ignored manipulation of the driver. To date, only one activation domain has been demonstrated to function efficiently in single copy to drive transcription. VP16, VP64 (4X repeat of VP16) and VPR (combination a VP64, NF-κB p65 subunit activation domain and Rta activator from Epstein-Barr Virus) all perform poorly in *C. elegans* (Mao et al., 2019), and only the activation domain of QF shows strong activity. It remains possible that a significantly more potent activator might be found for *C. elegans*. Identifying new activators is particularly important since QF can be regulated by expression of QS. If all bipartite systems use QF, one will not be able to regulate a system using QS independently of another.

### Basal promoters

As we have outlined above, expression of a reporter is clearly modulated by the number of driver binding sites present in a promoter. Additionally, basal promoters significantly influence transgene expression, with the maximal strength of expression limited by the ability of RNA Polymerase II to initiate at the promoter. Thus, defining strong basal promoters that have limited background activity is paramount to developing a robust and specific expression system. Little experimental work on basal promoters has been performed in *C. elegans* in the last decade. Several basal promoters were defined in the 1990’s and early 20’s and functionally they were described as ∼100-150 bp sequences upstream of the ATG in the *myo-2*, *hsp-16*, *mec-7*, and *pes-10* promoters (Okkema et al., 1993; Okkema and Fire, 1994; Duggan et al., 1998; Thatcher et al., 2001; Hong et al., 2004). More recently, genomic analysis has defined the molecular features of basal promoters (Grishkevich et al., 2011; Gu et al., 2012; Chen et al., 2013; Kruesi et al., 2013; Reinke et al., 2013; Saito et al., 2013). These analyses revealed that basal promoters are relatively heterogenous with multiple distinct motifs enriched proximal to the initiation site that are observed in a small fraction of individual promoters. Consistent with the bioinformatic analysis, we found that all the random 200-400 bp sequences tested can promote moderate levels of transcription in front of a *tetO* 7X enhancer. In a small survey of bona fide basal promoters, we focused on basal promoters of genes that either express very abundant cellular proteins or promoters that are highly inducible. Although, some of the basal promoter fragments identified express at high levels, none were free of background expression. The levels of background expression are very modest and are unlikely to hamper the use of these promoters for creating transgenes that aim to express reporters or gene products in select cell types at high levels. However, these promoters likely cannot confidently be used to define the cell-specific functions of a gene. The background leakiness of these promoters will require stringent controls for such analysis. Unexpectedly, some of the basal promoter characterized show relatively strong cell-specificity in their activity level. For example, the *mec-7* basal promoter is more robust in TRNs than the intestine, and the *tbb-2* basal promoter is more robust in the intestine than in TRNs. Since *mec-7* encodes a tubulin expressed selectively in TRNs, this may suggest that the defined basal promoter still contains cell specific regulatory sequences that either potentiate or repress transcription in a cell specific manner. In summary, an ideal basal promoter for creating reporter construct is still lacking, and further characterization of basal promoters is clearly warranted.

### 3’ UTRs

3 ‘UTRs have been previously documented to have powerful influence on expression levels (Merritt et al., 2008). One major mechanism of this control is through regulating the stability and translation of the mRNA via miRNAs (Ambros and Ruvkun, 2018). However, in bipartite reporter expression assays, we noticed common background signals that suggested that the 3’ UTRs might also be influencing transcription as enhancers. Several commonly used 3’ UTRs, including *tbb-2*, show such activity in enhancer assays developed to test this hypothesis. The ability to create many very similar transgenes at the same locus provides a straightforward assay to deduce which sequences exhibit such activity.

In addition to determining many facets of message stability and translation efficiency, the 3’ UTR region, including the sequences downstream of the poly adenylation signal, are also involved in transcriptional termination. My lab had previously presumed that 3’ UTRs terminated transcription efficiently. However, in building many transgenes that contain multiple closely spaced ‘independent’ transcription units, it has become clear that adjacent transcription units cannot simply be treated as independent elements. Cross transcription unit interactions at the level of transcription initiation are clearly contributing to unexpected expression observed in transgenes. However, inefficient transcriptional termination also has great potential to mediate cross transcription unit interactions. Furthermore, since multi-cistronic operons are common in *C. elegans* (Blumenthal and Gleason, 2003), the transcription apparatus has a built-in set of mechanisms to purposefully prevent transcriptional termination. In an assay designed to assess the efficiency of transcriptional termination, we observed large differences in the efficiency of the process. These data indicate that the choice of the 3’ UTR, the orientation of neighboring transcription units, and the spacing of these units are all likely to influence expression. While these studies only examined the influences on the components of our transgenic inserts, the inserted sequences could and likely will influence the expression of the endogenous *C. elegans* genes adjacent to the site of insertion, especially if poorly terminating 3’ UTRs are utilized.

### Direction of transcription

Textbook descriptions of transcription present it as a direction selective process. However, genome wide analyses of RNA species have shown that a large fraction of promoters initiate transcription in both directions (Seila et al., 2009) including in *C. elegans* (Gu et al., 2012; Chen et al., 2013; Kruesi et al., 2013; Saito et al., 2013). In direct tests for bi-directional transcription from a *tetO* 7X promoter, we observed strong expression of the reverse strand reporter. This effect was observed in the absence of a second basal promoter. Since, the *mec-7* basal promoter is required for forward transcription (Fig 6A), this suggests that the basal promoter directs the bi-directional transcription. Formally, we cannot exclude the possibility that initiation is also occurring from within the *tetO* operator sequence in a reverse direction specific manor. Regardless of the mechanism of this reverse transcription, it should be considered in the design of transgenes. One approach could be to introduce a 3’ UTR/terminator upstream of the *tetO* enhancer to rapidly terminate any reverse transcription.

### Influence of distance on expression

In designing a transgene, one usually presumes that reporter (e.g. *tetO*) enhancer functions by directing transcription at the juxtaposed basal promoter. However, in theory this enhancer could engage other neighboring promoters. In mammalian systems one mechanism to direct and control such interactions is inserting insulator elements, which act to physically segregate DNA regions to reduce crosstalk. However, no such insulator elements have been functionally defined compact *C. elegans* genome. Furthermore, the genes encoding the proteins which bind to classical insulator elements are absent from *C. elegans* (Heger et al., 2009). The absence of such elements eliminates an obvious tool to insulate transcription units in transgenes. To assess the properties of enhancer promoter interactions, we examined how distance between the *tetO* enhancer and the basal promoter elements influences expression. Distance was, in general, a strong negative influence on transcription. However, one of the transgenes evaded such distance effects suggesting that genome structural elements can strongly influence expression. Further study is clearly required to understand how the activity of regulatory elements is limited to specific promoters in the context of the closely spaced genome organization found in *C. elegans* and many other metazoans.

## Methods

### *C. elegans* strain maintenance

*C. elegans* was grown on NGM plates seeded with *E. coli* OP50 on 6 cm plates. Stock strains were maintained at RT (∼ 22.5°C). RMCE and rRMCE experiments were performed at 25°C.

### Plasmid and vector constructions

Plasmids were constructed either using Golden Gate reactions (Engler et al., 2008) or PCR followed by recircularization using T4 DNA ligase. All PCR amplification for plasmid and vector constructions were performed using Q5 polymerase (New England Biolabs, Ipswich, MA). Most PCR reactions were performed using the following conditions: 98°C for 0:30, followed by 30 cycles of 98°C for 0:10, 62°C for 0:30, 72°C for 1:00/kb). PCR products were digested with *DpnI* to remove template if amplified from a plasmid, then purified using a standard Monarch (New England Biolabs) column purification procedure. Restriction enzymes (except for *LguI*), T4 DNA ligase, and polynucleotide kinase were purchased from New England Biolabs. Golden Gate (GG) reactions (Engler et al., 2008) were performed as described in Nonet (2020) except that in some cases *LguI* (Thermo Scientific™, Waltam, MA) was used in place of *SapI*. The *E. coli* strain DH5α was used for all transformations. Sanger sequencing was performed by GENEWIZ (South Plainfield, NJ) and nanopore sequencing by Plasmidsaurus (Eugene, OR). Oligonucleotides were obtained from MilliporeSigma (Burlington, MA) and synthetic DNA fragment were purchased from Twist Biosciences (South San Francisco, CA). Many plasmids used in this study were recently described as part of a toolkit for assembly RMCE targeting clones (Knoebel et al., 2023). A detailed description of all constructs is provided in supplemental methods. The sequence of all plasmids, vectors and synthetic fragments and oligonucleotides used in the study are provided in Table S1.

### Creation of transgenic animals by microinjection

Injections were performed as described in Nonet (2020) except that a paint brush (Robert Simmons E51 liner 10/0) was typically used to mount animals onto the agar pad before injections. Most animals were only injected in a single gonad. DNAs were injected at ∼ 50 µg/ml in 10 mM Tris pH 8.0, 0.1 mM EDTA. Both RMCE and rRMCE were used to create transgenic insertions. RMCE insertions were performed as detailed in Nonet (2020) and rRMCE insertions as detailed in Nonet (2023). In some cases, the *cis* marker introduced during rRMCE was removed by outcrosses to germline Cre expressing strains Nonet (2023). Table S1 contains a list of all transgenic animals created in thus study and includes the plasmid used as a template, the *C. elegans* strain injected, and the sequence of the insertion. The molecular structure of transgenic insertions created using (r)RMCE is reliably identical to the predicted recombination product (Nonet, 2023). A small subset of insertions were molecular confirmed using either PCR (outlined in Nonet, 2020) or nanopore sequencing as listed in Table S1. However, the vast majority were not molecular characterized and are presumed to be correct based on the efficacy of RMCE.

### Microscopy

Screening of worms for fluorescence during analysis was performed on a Leica (Heerbrugg, Switzerland) MZ16F FluoCombi III microscope with a plan Apo 1X and plan Apo 5X LWD objective for high power observation illuminated using a Lumencor (Beaverton, OR) Sola light source.

For imaging or quantification of fluorescence, worms were mounted on 2 % agarose pads in a 2 µl drop of 1 mM levamisole in phosphate buffered saline. 10 to 20 L4 animals were typically placed on a single slide. Animals were imaged using a 10X air (n.a. 0.45) or 40X air (n.a. 0.75) lens on an Olympus (Center Valley, PA) BX-60 microscope. Initially images were obtained using a Qimaging (Surrey, BC Canada) Retiga EXi monochrome CCD camera, a Lumencor AURA LED light source, Semrock (Rochester, NY) GFP-3035B and mCherry-A-000 filter sets, a Chroma (Bellows Falls, VT) 89402 multi-pass filter set and a Tofra (Palo Alto, CA) focus drive, run using Micro-Manager 2.0ß software (Edelstein et al., 2014). However, due to equipment failure, later images were taken after replacing the Retiga with a ToupTek MAX04BM sCMOS camera. All data presented within individual figure panels used the same camera for acquisition. Images were quantified using the FIJI version of ImageJ software (Schindelin et al., 2012) as described in Nonet (2020). In cases where images were rotated leading to the presence of black corners in some panels, these black corners were adjusted to the mean intensity of a 3-pixel width line adjacent to the diagonal of each corner using a custom imageJ macro. Data plots were created using PlotsOfData (Postma and Goedhart, 2019).

## Data Availability

A full description of all oligonucleotides, plasmids, transgenes, and *C. elegans* strains created and used in this article are in Table S1. *C. elegans* strains to be sent to the Caenorhabditis Genetics Center are listed in Table S1. All other reagents are available upon request to Michael Nonet.

## Supplemental Materials

The manuscript contains 7 supplemental figures, one supplemental table and two supplemental text files named supplemental methods and supplemental figure legends.

## Acknowledgements

I thank Dany Matar for creating several driver clones, Jordan Ward, Matt Rich, and Han Wang and especially Tim Schedl for valued discussions. Some strains were provided by the CGC, which is funded by NIH Office of Research Infrastructure Programs (P40 OD010440).

## Funding

This work was supported in part by the National Institutes of Health grant R01GM14168802 awarded to MLN.

## Conflicts of interest

The authors declare no conflict of interest.

## Abbreviations

TRN: touch receptor neuron
RMCE: Recombination-Mediated Cassette Exchange
rRMCE: rapid RMCE
3’ UTR: three prime untranslated region
mNG: monomeric Neon Green FP
FP: Fluorescent Protein
GFP: Green FP
SEC: Self-Excision Cassette

## References

Agha-Mohammadi S, O’malley M, Etemad A, Wang Z, Xiao X, Lotze MT. Second-generation tetracycline-regulatable promoter: repositioned tet operator elements optimize transactivator synergy while shorter minimal promoter offers tight basal leakiness. J Gene Med. 2004;6(7):817–828.

Ahringer J, Gasser SM. Repressive Chromatin in Caenorhabditis elegans: Establishment, Composition, and Function. Genetics. 2018;208(2):491–511.

Akay A, Jordan D, Navarro IC, Wrzesinski T, Ponting CP, Miska EA, Haerty W. Identification of functional long non-coding RNAs in C. elegans. BMC Biology. 2019;17(1):; doi: 10.1186/s12915-019-0635-7.

Ambros V, Ruvkun G. Recent Molecular Genetic Explorations of Caenorhabditis elegans MicroRNAs. Genetics. 2018;209(3):651–673.

Arribere JA, Bell RT, Fu BX, Artiles KL, Hartman PS, Fire AZ. Efficient Marker-Free Recovery of Custom Genetic Modifications with CRISPR/Cas9 in Caenorhabditis elegans. Genetics. 2014; doi: 10.1534/genetics.114.169730.

Artan M, Barratt S, Flynn SM, Begum F, Skehel M, Nicolas A, De Bono M. Interactome analysis of Caenorhabditis elegans synapses by TurboID-based proximity labeling. J Biol Chem. 2021;297(3):101094.

Bagchi DN, Lyer VR. The Determinants of Directionality in Transcriptional Initiation. Trends Genet. 2016;32(6):322–333.

Bakowski MA, Desjardins CA, Smelkinson MG, Dunbar TL, Dunbar TA, Lopez-Moyado IF, Rifkin SA, Cuomo CA, Troemel ER. Ubiquitin-mediated response to microsporidia and virus infection in C. elegans. PLoS Pathog. 2014;10(6):e1004200.

Blumenthal T, Gleason KS. Caenorhabditis elegans operons: form and function. Nat Rev Genet. 2003;4(2):112–120.

Brand AH, Perrimon N. Targeted gene expression as a means of altering cell fates and generating dominant phenotypes. Development. 1993;118(2):401–415.

Cao J, Packer JS, Ramani V, Cusanovich DA, Huynh C, Daza R, Qiu X, Lee C, Furlan SN, Steemers FJ, Adey A, Waterston RH, Trapnell C, Shendure J. Comprehensive single-cell transcriptional profiling of a multicellular organism. Science. 2017;357(6352):661–667.

Caygill EE, Brand AH. The GAL4 System: A Versatile System for the Manipulation and Analysis of Gene Expression. Methods Mol Biol. 2016;1478:33–52.

Chai Y, Li W, Feng G, Yang Y, Wang X, Ou G. Live imaging of cellular dynamics during Caenorhabditis elegans postembryonic development. Nat Protoc. 2012;7(12):2090–2102.

Chen K, Franz CJ, Jiang H, Jiang Y, Wang D. An evolutionarily conserved transcriptional response to viral infection in Caenorhabditis nematodes. BMC Genomics. 2017;18(1):303.

Chen RA-J, Down TA, Stempor P, Chen QB, Egelhofer TA, Hillier LW, Jeffers TE, Ahringer J. The landscape of RNA polymerase II transcription initiation in C. elegans reveals promoter and enhancer architectures. Genome Research. 2013;23(8):1339–1347.

Davis P, Zarowiecki M, Arnaboldi V, Becerra A, Cain S, Chan J, Chen WJ, Cho J, Da Veiga Beltrame E, Diamantakis S, Gao S, Grigoriadis D, Grove CA, Harris TW, Kishore R, Le T, Lee RYN, Luypaert M, Müller HM, Nakamura C, Nuin P, Paulini M, Quinton-Tulloch M, Raciti D, Rodgers FH, Russell M, Schindelman G, Singh A, Stickland T, Van Auken K, Wang Q, Williams G, Wright AJ, Yook K, Berriman M, Howe KL, Schedl T, Stein L, Sternberg PW. WormBase in 2022-data, processes, and tools for analyzing Caenorhabditis elegans. Genetics. 2022;220(4):iyac003.

Dickinson DJ, Ward JD, Reiner DJ, Goldstein B. Engineering the Caenorhabditis elegans genome using Cas9-triggered homologous recombination. Nat Methods. 2013;10:1028–1034.

Distel M, Wullimann MF, Köster RW. Optimized Gal4 genetics for permanent gene expression mapping in zebrafish. Proc Natl Acad Sci U S A. 2009;106(32):13365–13370.

Dour S, Nonet M. Optimizing expression of a single copy transgene in C. elegans. MicroPubl Biol. 2021;2021:10.17912/micropub.biology.000394.

Duggan A, Ma C, Chalfie M. Regulation of touch receptor differentiation by the Caenorhabditis elegans mec-3 and unc-86 genes. Development. 1998;125(20):4107–4119.

Edelstein AD, Tsuchida MA, Amodaj N, Pinkard H, Vale RD, Stuurman N. Advanced methods of microscope control using μManager software. J Biol Methods. 2014;1(2):e10.

El Mouridi S, Alkhaldi F, Frøkjær-Jensen C. Modular safe-harbor transgene insertion for targeted single-copy and extrachromosomal array integration in Caenorhabditis elegans. G3 (Bethesda). 2022;12(9):jkac184.

Engler C, Kandzia R, Marillonnet S. A one pot, one step, precision cloning method with high throughput capability. PLoS One. 2008;3(11):e3647.

Fashena SJ, Serebriiskii IG, Golemis EA. LexA-based two-hybrid systems. Methods Enzymol. 2000;328:14–26.

Fire A, Alcazar R, Tan F. Unusual DNA structures associated with germline genetic activity in Caenorhabditis elegans. Genetics. 2006;173(3):1259–1273.

Frokjaer-Jensen C, Davis MW, Hopkins CE, Newman BJ, Thummel JM, Olesen SP, Grunnet M, Jorgensen EM. Single-copy insertion of transgenes in Caenorhabditis elegans. Nat Genet. 2008;40(11):1375–1383.

Frøkjær-Jensen C, Davis MW, Sarov M, Taylor J, Flibotte S, Labella M, Pozniakovsky A, Moerman DG, Jorgensen EM. Random and targeted transgene insertion in Caenorhabditis elegans using a modified Mos1 transposon. Nat Methods. 2014;11(5):529–534.

Gossen M, Bujard H. Tight control of gene expression in mammalian cells by tetracycline-responsive promoters. Proc Natl Acad Sci U S A. 1992;89(12):5547–5551.

Graber JH, Salisbury J, Hutchins LN, Blumenthal T. C. elegans sequences that control trans-splicing and operon pre-mRNA processing. RNA. 2007;13(9):1409–1426.

Grishkevich V, Hashimshony T, Yanai I. Core promoter T-blocks correlate with gene expression levels in C. elegans. Genome Res. 2011;21(5):707–717.

Gu W, Lee HC, Chaves D, Youngman EM, Pazour GJ, Conte D, Mello CC. CapSeq and CIP-TAP identify Pol II start sites and reveal capped small RNAs as C. elegans piRNA precursors. Cell. 2012;151(7):1488–1500.

Haenni S, Sharpe HE, Gravato Nobre M, Zechner K, Browne C, Hodgkin J, Furger A. Regulation of transcription termination in the nematode Caenorhabditis elegans. Nucleic Acids Res. 2009;37(20):6723–6736.

Heger P, Marin B, Schierenberg E. Loss of the insulator protein CTCF during nematode evolution. BMC Mol Biol. 2009;10:84–97.

Hong M, Kwon JY, Shim J, Lee J. Differential hypoxia response of hsp-16 genes in the nematode. J Mol Biol. 2004;344(2):369–381.

Knoebel E, Dour S, Nonet ML. A toolkit for assembly of targeting clones for C. elegans transgenesis. MicroPubl Biol. 2023;2023:10.17912/micropub.biology.000966.

Kruesi WS, Core LJ, Waters CT, Lis JT, Meyer BJ. Condensin controls recruitment of RNA polymerase II to achieve nematode X-chromosome dosage compensation. Elife. 2013;2:e00808.

Li W, Notani D, Rosenfeld MG. Enhancers as non-coding RNA transcription units: recent insights and future perspectives. Nat Rev Genet. 2016;17(4):207–223.

Mahoney TR, Luo S, Round EK, Brauner M, Gottschalk A, Thomas JH, Nonet ML. Intestinal signaling to GABAergic neurons regulates a rhythmic behavior in Caenorhabditis elegans. Proc Natl Acad Sci U S A. 2008;105(42):16350–16355.

Mao S, Qi Y, Zhu H, Huang X, Zou Y, Chi T. A Tet/Q Hybrid System for Robust and Versatile Control of Transgene Expression in C. elegans. iScience. 2019;11:224–237.

Merritt C, Rasoloson D, Ko D, Seydoux G. 3’ UTRs are the primary regulators of gene expression in the C. elegans germline. Curr Biol. 2008;18(19):1476–1482.

Miki TS, Carl SH, Großhans H. Two distinct transcription termination modes dictated by promoters. Genes Dev. 2017;31(18):1870–1879.

Nadiminti SSP, Koushika SP. Imaging Intracellular Trafficking in Neurons of C. elegans. Methods Mol Biol. 2022;2431:499–530.

Nance J, Frøkjær-Jensen C. The Caenorhabditis elegans Transgenic Toolbox. Genetics. 2019;212(4):959–990.

Noma K, Jin Y. Rapid Integration of Multi-copy Transgenes Using Optogenetic Mutagenesis in Caenorhabditis elegans. G3 (Bethesda). 2018;8(6):2091–2097.

Nonet M. Improved GAL4 and Tet OFF drivers for C. elegans bipartite expression. MicroPubl Biol. 2021;2021:10.17912/micropub.biology.000438.

Nonet ML. Efficient transgenesis in C. elegans using Flp recombinase-mediated cassette exchange. Genetics. 2020;215:902–921.

Nonet ML. Rapid generation of C. elegans single-copy transgenes combining RMCE and drug selection. Genetics. 2023 iyad072.

Okkema PG, Harrison SW, Plunger V, Aryana A, Fire A. Sequence requirements for myosin gene expression and regulation in Caenorhabditis elegans. Genetics. 1993;135(2):385–404.

Okkema PG, Fire A. The Caenorhabditis elegans NK-2 class homeoprotein CEH-22 is involved in combinatorial activation of gene expression in pharyngeal muscle. Development. 1994;120(8):2175–2186.

Paix A, Wang Y, Smith H, Lee C-YS, Calidas D, Lu T, Smith J, Schmidt H, Krause M, Seydoux G. Scalable and Versatile Genome Editing Using Linear DNAs with Micro-homology to Cas9 Sites in Caenorhabditis elegans. Genetics. 2014; doi: 10.1534/genetics.114.170423.

Pfeiffer BD, Ngo T-TB, Hibbard KL, Murphy C, Jenett A, Truman JW, Rubin GM. Refinement of tools for targeted gene expression in Drosophila. Genetics. 2010;186(2):735–755.

Postma M, Goedhart J. PlotsOfData-A web app for visualizing data together with their summaries. PLoS Biol. 2019;17(3):e3000202.

Potter CJ, Tasic B, Russler EV, Liang L, Luo L. The Q system: a repressible binary system for transgene expression, lineage tracing, and mosaic analysis. Cell. 2010;141(3):536–548.

Quintero-Cadena P, Sternberg PW. Enhancer Sharing Promotes Neighborhoods of Transcriptional Regulation Across Eukaryotes. G3 (Bethesda). 2016;6(12):4167–4174.

Reinke V, Krause M, Okkema P. Transcriptional regulation of gene expression in C. elegans. WormBook. 2013 1–34.

Riabinina O, Potter CJ. The Q-System: A Versatile Expression System for Drosophila. Methods Mol Biol. 2016;1478:53–78.

Saito TL, Hashimoto S, Gu SG, Morton JJ, Stadler M, Blumenthal T, Fire A, Morishita S. The transcription start site landscape of C. elegans. Genome Res. 2013;23(8):1348–1361.

Schindelin J, Arganda-Carreras I, Frise E, Kaynig V, Longair M, Pietzsch T, Preibisch S, Rueden C, Saalfeld S, Schmid B, Tinevez JY, White DJ, Hartenstein V, Eliceiri K, Tomancak P, Cardona A. Fiji: an open-source platform for biological-image analysis. Nat Methods. 2012;9(7):676–682.

Schönig K, Bujard H, Gossen M. The power of reversibility regulating gene activities via tetracycline-controlled transcription. Methods Enzymol. 2010;477:429–453.

Schwartz ML, Davis MW, Rich MS, Jorgensen EM. High-efficiency CRISPR gene editing in C. elegans using Cas9 integrated into the genome. 2021; doi: 10.1101/2021.08.03.454883.

Seila AC, Core LJ, Lis JT, Sharp PA. Divergent transcription: a new feature of active promoters. Cell Cycle. 2009;8(16):2557–2564.

Silva-García CG, Lanjuin A, Heintz C, Dutta S, Clark NM, Mair WB. Single-Copy Knock-In Loci for Defined Gene Expression in Caenorhabditis elegans. G3 (Bethesda). 2019;9(7):2195–2198.

Stevenson ZC, Moerdyk-Schauwecker MJ, Banse SA, Patel DS, Lu H, Phillips PC. High-throughput library transgenesis in Caenorhabditis elegans via Transgenic Arrays Resulting in Diversity of Integrated Sequences (TARDIS). Elife. 2023;12:RP84831.

Thatcher JD, Fernandez AP, Beaster-Jones L, Haun C, Okkema PG. The Caenorhabditis elegans peb-1 gene encodes a novel DNA-binding protein involved in morphogenesis of the pharynx, vulva, and hindgut. Dev Biol. 2001;229(2):480–493.

Thomas-Chollier M, Defrance M, Medina-Rivera A, Sand O, Herrmann C, Thieffry D, Van Helden J. RSAT 2011: regulatory sequence analysis tools. Nucleic Acids Res. 2011;39(Web Server issue):W86–91.

Towbin BD, González-Aguilera C, Sack R, Gaidatzis D, Kalck V, Meister P, Askjaer P, Gasser SM. Step-wise methylation of histone H3K9 positions heterochromatin at the nuclear periphery. Cell. 2012;150(5):934–947.

Venkatachalam V, Ji N, Wang X, Clark C, Mitchell JK, Klein M, Tabone CJ, Florman J, Ji H, Greenwood J, Chisholm AD, Srinivasan J, Alkema M, Zhen M, Samuel AD. Pan-neuronal imaging in roaming Caenorhabditis elegans. Proc Natl Acad Sci U S A. 2016;113(8):E1082–8.

Wang H, Liu J, Gharib S, Chai CM, Schwarz EM, Pokala N, Sternberg PW. cGAL, a temperature-robust GAL4-UAS system for Caenorhabditis elegans. Nat Methods. 2017;14(2):145–148.

Wei X, Potter CJ, Luo L, Shen K. Controlling gene expression with the Q repressible binary expression system in Caenorhabditis elegans. Nat Methods. 2012;9(4):391–395.

Yagi R, Mayer F, Basler K. Refined LexA transactivators and their use in combination with the Drosophila Gal4 system. Proceedings of the National Academy of Sciences. 2010;107(37):16166–16171.

Yang FJ, Chen CN, Chang T, Cheng TW, Chang NC, Kao CY, Lee CC, Huang YC, Hsu JC, Li J, Lu MJ, Chan SP, Wang J. phiC31 integrase for recombination-mediated single-copy insertion and genome manipulation in Caenorhabditis elegans. Genetics. 2022;220(2):iyab206.

Yoshina S, Suehiro Y, Kage-Nakadai E, Mitani S. Locus-specific integration of extrachromosomal transgenes in C. elegans with the CRISPR/Cas9 system. Biochem Biophys Rep. 2016;5:70–76.

Zhang L, Ward JD, Cheng Z, Dernburg AF. The auxin-inducible degradation (AID) system enables versatile conditional protein depletion in C. elegans. Development. 2015;142(24):4374–4384.

